# Informed movement and dispersal in experimental metacommunities

**DOI:** 10.1101/017954

**Authors:** Emanuel A. Fronhofer, Jan Klecka, Carlos J. Melián, Florian Altermatt

**Affiliations:** Eawag: Swiss Federal Institute of Aquatic Science and Technology, Department of Aquatic Ecology, Überlandstrasse 133, CH-8600 Dübendorf, Switzerland; Institute of Evolutionary Biology and Environmental Studies, University of Zurich, Winterthurerstrasse 190, CH-8057 Zürich, Switzerland; Eawag: Swiss Federal Institute of Aquatic Science and Technology, Department of Fish Ecology and Evolution, Seestrasse 79, CH-6047 Kastanienbaum, Switzerland; Laboratory of Theoretical Ecology, Institute of Entomology, Biology Centre of the Czech Academy of Sciences, České Budějovice, Czech Republic

**Keywords:** inter-specific competition, plasticity, reaction norm, density-dependent dispersal, density-dependent movement, Allee effect, microcosms, protists

## Abstract

Dispersal, and the underlying movement behaviour, are processes of pivotal importance for understanding and predicting metapopulation and metacommunity dynamics. Generally, dispersal decisions are non-random and rely on information, such as the presence of conspecifics. However, studies on metacommunities that include interspecific interactions generally disregard information use. Therefore, it remains unclear whether and how dispersal in metacommunities is informed and whether rules derived from single-species contexts can be scaled up to (meta-)communities. Using experimental protist metacommunities, we show how dispersal and movement are informed and adjusted by the strength of inter-specific interactions. We found that predicting informed movement and dispersal in metacommunities requires knowledge on behavioural responses to intra- and inter-specific interaction strength. Consequently, metacommunity dynamics inferred directly from single-species metapopulations without taking inter-specific interactions into account are likely flawed. Our work identifies the significance of information use for understanding metacommunity dynamics, stability and the coexistence and distribution of species.

**Author contributions:** All authors designed the study. EAF and JK performed the experiments and analysed the data. EAF wrote the manuscript and all authors contributed substantially to revisions.

## Introduction

Local populations and communities are usually not isolated entities, but rather part of a larger spatially structured system (Hanski, 1999; Leibold *et al.*, 2004). Therefore, dispersal of individuals between these local communities is by definition a process of central relevance for both local and global ecological and evolutionary dynamics (Clobert *et al.*, 2012). Dispersal in such metapopulations and metacommunities has been intensely studied over the last decades and it is now clear that dispersal and movement can be highly complex behaviours (Nathan *et al.*, 2008; Clobert *et al.*, 2012) with important consequences, even for large-scale patterns such as metapopulation dynamics (Fronhofer *et al.*, 2012), species range dynamics (Kubisch *et al.*, 2014), species coexistence (Salomon *et al.*, 2010), or the distribution of biological diversity (Seymour *et al.*, 2015).

Both empirical and theoretical studies show that dispersal as well as the underlying movement behaviour are fundamentally non-random processes (Nathan *et al.*, 2008; Clobert *et al.*, 2009). Information use is highly advantageous during all three stages of dispersal (emigration, transition, immigration) and a variety of cues, such as local population density (Matthysen, 2005; De Meester & Bonte, 2010; Bitume *et al.*, 2013; Fronhofer *et al.*, 2015), food availability and autocorrelation (Kuefler *et al.*, 2012; Fronhofer *et al.*, 2013a), relatedness (Bitume *et al.*, 2013), body condition (Bonte & de la Peña, 2009), chemical cues (Fronhofer *et al.*, 2013b) or abiotic conditions (Crone *et al.*, 2001; Altermatt & Ebert, 2010; Reigada *et al.*, 2015) can be used to trigger dispersal and movement decisions. Therefore, dispersal and movement are generally condition-dependent or informed and not fixed traits.

Informed dispersal and movement can have far-reaching consequences such as altered (macro-)ecological and evolutionary dynamics. For instance, source-sink dynamics were shown to be impacted by density-dependent dispersal (Amarasekare, 2004) and species ranges are predicted to be larger (Kubisch *et al.*, 2011) and invasions slower (Altwegg *et al.*, 2013) when dispersal is positively density-dependent. By contrast, negatively density-dependent dispersal may increase invasion speeds (Altwegg *et al.*, 2013). Finally, evolutionarily stable dispersal rates are usually higher for non-conditional dispersers implying higher costs (Enfjäll & Leimar, 2009), to name just a few consequences.

Most studies on informed dispersal have focused on the effects of intra-specific density (Matthysen, 2005; De Meester & Bonte, 2010; Bitume *et al.*, 2013; Fronhofer *et al.*, 2015), largely ignoring effects of species interactions and inter-specific densities. However, species rarely exist in isolation and, in a metacommunity context, both dispersal and movement should also depend on inter-specific interactions. However, metacommunity studies usually assume that dispersal is a random process that does not depend on metacommunity composition (but see Amarasekare 2010 for a theoretical model and De Meester *et al.* 2015 for empirical evidence). Clearly, this is a relatively unlikely assumption, and for the specific cases of host-parasitoid (French & Travis, 2001) and predator-prey systems (Poethke *et al.*, 2010; Hauzy *et al.*, 2007; Kuefler *et al.*, 2012) it has been shown that movement and dispersal are condition-dependent and modulated by the abundance of the antagonist. Surprisingly, the effect of omnipresent competitive interactions on movement and dispersal have been ignored. Therefore, we are lacking an understanding of the effects of competition among species of the same trophic level on dispersal and movement. Thus, it remains unclear whether one can simply infer these effects from intra-specific density-dependence or if inter-specific interactions modulate movement and dispersal dynamics. Such knowledge, however, is crucial and a prerequisite for building a reliable, predictive science of ecological dynamics in space.

We tested experimentally whether and how one can generalize models of informed, density-dependent dispersal and movement, which have been developed for single-species metapopulations, to be applicable to a multi-species, metacommunity context. Intra-specific density-dependent dispersal has been shown theoretically to be generally positive as a means to escape from competition, with the exact shape of the function depending on model assumptions (Metz & Gyllenberg, 2001; Poethke & Hovestadt, 2002). Negative density-dependence (see e.g. Matthysen, 2005; Baguette *et al.*, 2011; Fellous *et al.*, 2012) may arise for example at low population densities due to Allee effects (Kim *et al.*, 2009; Fronhofer *et al.*, 2015), which, in a single-species framework, is predicted to lead to a u-shaped density-dependent dispersal and movement reaction norm Fig. 1). In a multi-species context we predict that dispersal and movement fundamentally follow the same u-shaped relationship but that conspecific and all heterospecific densities weighted by their respective interaction strength have to be taken into account to explain and predict dispersal and movement behaviour correctly (see Fig. 1 for details). To test our prediction in a multispecies context we focused on metacommunities of competitors, i.e. species of the same trophic level, reflecting some of the most-studied and most relevant interactions (e.g. Chesson, 2000; Hubbell, 2001). Based on experimentally assessed differences in competition strength we identified the effects of competition on movement and dispersal across levels of community diversity form single-species to all two- and three-species metacommunites of three protist model organisms (Altermatt *et al.*, 2015). Specifically, we answer the following questions: (1) Are movement and dispersal informed with regards to inter-specific competition in metacommunities as one would predict in analogy to intra-specific density-dependent dispersal (Fig. 1)? (2) Can the patterns observed in simple two-species metacommunities be scaled up to predict the behaviour of species in a more diverse, three-species metacommunity?

**Figure 1:**
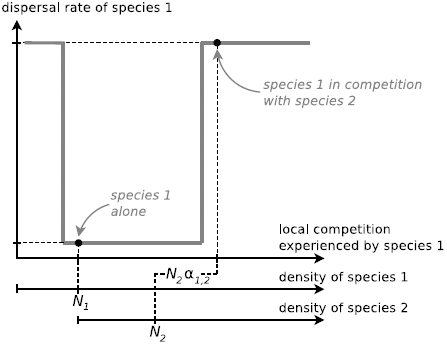
Predictions about the impact of inter-specific competition on informed dispersal and movement in a metacommunity consisting of species experiencing an Allee effect. In a single species context, one can expect species that suffer from an Allee effect to exhibit roughly u-shaped density-dependent dispersal and movement functions (reaction norm; grey line). Conspecific density will then determine the dispersal rate or movement strategy (e.g., *d*_*N*_1__, “species 1 alone”). If this model is generally valid and can be transferred to the metacommunity context, the dispersal rate and movement strategy of the focal species (species 1) should be determined by the sum of its own density (*N*_1_) and the density of competitors weighted by the respective competition coefficient (*N*_2_ *α*_1,2_) and result in an altered dispersal/ movement strategy (e.g., *d*_*N*_1_*with N*_2__, “species 1 in competition with species 2”).

Our results show that movement and dispersal are informed with regards to inter-specific competition. Specifically, inter-specific interaction strength and u-shaped density-dependent dispersal and movement reaction norms (Fig. 1) in combination with knowledge about the behavioural response of competing species can be used to explain dispersal and movement behaviour across metacommunities. The ability to predict dispersal is a prerequisite for successfully scaling from simple, single-species to diverse and potentially complex multi-species systems and for understanding and predicting multispecies metacommunity dynamics and thus spatial dynamics in general.

## Materials and Methods

### Study organisms

We used a set of three freshwater protist species that are commonly used in experimental metacommunity studies: *Tetrahymena pyriformis* (Tet), *Colpidium* sp. (Col) and *Paramecium aurelia* (Pau). These ciliate species cover a wide range of growth rates (Pau *∼* Col *<* Tet; Altermatt *et al.* 2015), equilibrium densities (Pau *<* Col *<* Tet; Altermatt *et al.* 2015; see also Fig. S1 and Tab. S1) and body sizes (Tet *<* Col *<* Pau; Giometto *et al.* 2013; see also Supporting Information Fig. S3). Importantly, the three species differ in competitive abilities (Carrara *et al.* 2015; see also Tab. 1, Fig. S2 and Tab. S3). *Tetrahymena pyriformis* is known to suffer from an Allee effect (Chaine *et al.*, 2010; Fronhofer *et al.*, 2015), with implies reduced growth at low population densities. The underlying chemical mechanisms are general enough to allow the conclusion that both other species likely also have Allee effects which has been recently suggested for *Colpidium* and *Paramecium caudatum* (e.g. Odum & Allee, 1954; Duncan *et al.*, 2011). The presence of Allee effects in all three species studied is also supported by our data (see Tab. S2).

**Table 1:** Interaction coefficients (*α*_*i,j*_) of a Lotka-Volterra model of inter-specific competition. *α*_*i,j*_ represents the relative effect of species *j* on *i*. For example, if *α*_*i,j*_ = 2 species *j* has twice the competitive effect on species *i* as species *i* has on itself, *α*_*i,j*_ = 1 implies that inter- and intra-specific competition are equivalently strong and *α*_*i,j*_ = 0 suggests that species *j* does not compete with species *i*. Species *i* can be found in the columns and species *j* in the rows, so that *α*_*Tet,Col*_ = 2.85, for example. Reported values are median values of 8 independently fitted replicates. See Fig. S2 for community dynamics and model fits and Tab. S3 for information on variation.

|  | Tet | Col | Pau |
| --- | --- | --- | --- |
| Tet | 1 | 0.32 | 0.06 |
| Col | 2.85 | 1 | 0.01 |
| Pau | 18 | 2.53 | 1 |

Protist (meta)communities and single species (meta)populations were kept in protist medium (Protozoan pellets; Carolina Biological Supply, Burlington, USA; 0.46 g/L) at a constant temperature of 22*°* C (for detailed protocols see Altermatt *et al.*, 2015). Resources were supplied as 5% equilibrium density bacterial culture (approx. 1 week old; *Serratia fonticola*, *Bacillus subtilis* and *Brevibacillus brevis*) per litre of medium.

## Microcosm experimental design

We used two-patch microcosms for all experiments. In order to investigate the dependence of dispersal and movement on the presence of competing species we manipulated community composition in one patch (the start patch) and measured dispersal and movement to the second, initially empty patch (the target patch). These experimental, spatially structured systems consisted of two 20 mL vials (Sarstedt, Nümbrecht, Germany; distance between vials: 4 cm) which were connected by silicone tubes (inside diameter: 4 mm; VWR, Radnor, USA). Using clamps we controlled whether the connecting tubes were open or closed, i.e. dispersal could occur or not.

Before the experiment batch cultures of *Tetrahymena pyriformis*, *Colpidium* sp. and *Paramecium aurelia* were grown to carrying capacity (see Fig. S1). Subsequently, the batch cultures were centrifuged (centrifuge: Sigma 3-16PK; Sigma, Osterode, Germany; 4500 rpm; 5 min), the supernatant liquid was discarded and the pellet, which contained the protists, was resuspended in freshly bacterized medium (see Altermatt *et al.*, 2015). This washing procedure largely eliminated chemical cues that were present in the batch cultures. Such chemical cues were previously found to potentially trigger intra-specific density-dependent movement (Fronhofer *et al.*, 2015). We are therefore confident that the dispersal and movement behaviour observed here was not biased by cues remaining from the batch cultures. This implies that the differences in movement and dispersal between single-species metapopulations and multi-species metacommunities observed here can be attributed to an informed response with regards to inter-specific interactions since we controlled for intra-specific density.

Depending on the treatment (single-species metapopulations, two- or three-species metacommunities) start patches received populations or communities of one (Tet, Col or Pau), two (Tet–Col, Tet–Pau and Col–Pau) or three protist species (Tet–Col–Pau) from the washed batch cultures. In order to make consistent comparisons across all treatments individual species were always added at one fourth of their respective carrying capacity and filled up with freshly bacterized medium to a total volume of 15 mL. Target patches always received 15 mL freshly bacterized medium without protists.

Communities acclimatized to the new conditions in the start patch for one hour, during which protists could acquire information on abundance and presence of conspecifics and allospecifics. Then, the connecting tubes were opened and dispersal between the start and the target patch was allowed for 14 h. This period of time was determined trough pilot experiments in order to allow the least dispersive species (Pau) to be detected in the target patch (this was usually the case after approx. 10 to 12 h). After the dispersal phase, clamps were closed and movement as well as dispersal data was collected. All species combinations (treatments) were replicated 8 times in independent blocks.

## Data collection

### Movement

We collected individual-based movement (velocity, tortuosity, net distance travelled) and morphological data (length of the protist along major and minor axes, area) after the dispersal phase from the communities in the start patch using video analysis (for detiled protocols see Pennekamp *et al.*, 2014; Altermatt *et al.*, 2015). We followed the same procedure as described in Fronhofer *et al.* (2015): using a Nikon SMZ1500 stereo-microscope (Nikon Corporation, Kanagawa, Japan) with a Hamamatsu Orca Flash 4 video camera (Hamamatsu Photonics K.K., Hamamatsu city, Japan) we recorded videos for 20 s (500 frames) at a 20-fold magnification.

Subsequently, we used the free image analysis software ImageJ (version 1.46a U.S. National Institutes of Health, Bethesda, MD, USA, http://imagej.nih.gov/ij/) and the MOSAIC particle tracker plug-in (Sbalzarini & Koumoutsakos, 2005) for the video analysis. By sequentially subtracting frames from the video, the software determines moving particles of a given size range (particle area determined by previous experimentation: 5 to 600 pixels). Thereafter, the particle tracker plug-in re-links the locations taking into account a predefined link distance (here: 20 pixels) over subsequent frames (for details of the algorithm refer to Sbalzarini & Koumoutsakos, 2005). As described in Fronhofer *et al.* (2015) we used these movement paths to extract velocities, tortuosity (the circular standard deviation of the turning angle distribution) and net distance travelled (Euclidean distance from start to end of a movement path) as descriptive parameters that characterize a movement path.

Species within metacommunities were identified after the video analysis based on the collected morphological parameters (size and aspect ratio; see Fig. S3). We used a classification tree model (Statistical Software Package R version 3.1.2; package “tree” version 1.0-35, function “tree”) with aspect ratio and length along the major body axis as explanatory variables.

### Dispersal

Data on dispersal was collected by manually counting and identifying a representative sub-sample of protists from both start and target patches that had been conserved with Lugol after the experiment (for a detailed protocol see Altermatt *et al.*, 2015). As the species varied widely in densities (Fig. S1 and Tab. S1) sub-sample volumes and rates were adjusted accordingly (0.01 – 1 ml; 2 – 15 sub-samples). Dispersal rates were calculated as the ratio of dispersers (individuals per volume in the target patch) to the sum of all individuals (sum of individuals per volume in start and target patch) at the end of the experiment, i.e. after 14 h.

The time window we used for dispersal was relatively long, especially in relation to the reproductive rate of the fastest reproducing species (Tet; Fig. S1 and Tab. S1). Therefore, the measured dispersal rates could potentially be biased as a consequence of differential growth in start and target patch. In order to exclude this source of bias in our data we used a two-patch metapopulation model (Eqns. S4 – S7) which we fit using a least squares approach to the collected population density data from both patches. This model allowed us to estimate the corrected dispersal rates that take into account density-dependent growth in both patches and the potential impact of Allee effects (see above). While the corrected and measured dispersal rates differed from each other, these values correlated highly (LM: *N* = 23, *t* = 40.58, *p <* 0.001, *R*^2^ = 0.987; Fig. S5; for more detailed information and results, see Supporting Information). We therefore performed all analyses on the original, raw data.

### Population growth and inter-specific competition

The necessary data to estimate growth rates (*r*_0_), carrying capacities (*K*) as well as all pairwise inter-specific competition coefficients (*α*_*i,j*_; relative effect of species *j* on species *i*) were collected in separate experiments using the same microcosms and experimental setup, yet without allowing for dispersal. We subsequently fit logistic growth curves, respectively Lotka-Volterra models of inter-specific competition to the collected time-series data using a least-squares approach and extracted the relevant parameter estimates. See Supporting Information for details. Growth parameters are reported in the Tab. S1 and competition coefficients can be found in Tab. 1 and in more detail in Tab. S3.

## Statistical analysis

Data were analysed at the population level (means over all individuals) using linear mixed models (LMMs) with “replicate” as a random effect. This random effect structure was used as individual replicates including all species combinations were run in temporally independent blocks. In case of overdispersion an observation-level random effect was added. Specifically, dispersal rates were analysed as ratios of counts using generalized linear mixed models (GLMM) with a binomial error structure. In case other data did not allow a Gaussian error structure an appropriate GLMM was used. All statistical analyses were performed with the software package R version 3.1.2. (functions “lmer” and “glmer” from the “lmerTest” package; version 2.0-6). For pairwise comparisons we used Tukey post-hoc tests (function “glht” from the “multcomp” package; version 1.3-7).

The relative difference in movement and dispersal between single-species metapopulations and multispecies metacommunities was analysed using one-sample t-tests (*μ* = 0). If needed, data were transformed to satisfy the normality assumption. See Tab. S5 for further information.

## Results

### Relationship between movement and dispersal

We found large differences in dispersal rates between species: *Colpidium* sp. showed an order of magnitude higher dispersal rates compared to both *Tetrahymena pyriformis* and *Paramecium aurelia* Fig. 2 A ; see Tab. S4 for the detailed and full overview of all statistical analyses). Analogous differences were found in movement behaviour (Fig. 2B–D; Tab. S4): just as for dispersal, *Colpidium* sp. exhibited the highest net distance travelled. Generally, differences in net distance travelled were reflected in differences in velocity and tortuosity. Overall, higher velocities and less tortuous paths (narrower turning angle distribution) resulted in larger net distances travelled and vice versa (Fig. 2B–D; Tab. S4).

The qualitative correspondence of dispersal and movement reported for single-species metapopulations was found more generally, across all three species and in all metapopulations and -communities analysed in our experiments (Fig. S6). The strong non-linearity of the relationship between movement and dispersal is in good accordance with the results of Figs. 2: small differences in movement (Fig. 2B–D) potentially led to large differences in dispersal (Fig. 2 A).

**Figure 2:**
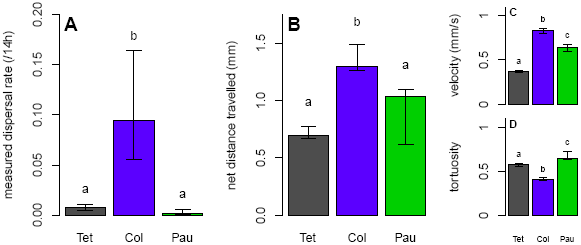
Dispersal and movement in single species metapopulations. (A) Measured dispersal rates for *Tetrahymena pyriformis* (Tet), *Colpidium* sp. (Col) and *Paramecium aurelia* (Pau) in single species, two-patch metapopulations. (B) Net distance travelled, i.e. distance from start to end of one individual movement trajectory in the same metapopulations. (C) Velocity. (D) Tortuosity measured as the standard deviation of the circular turning-angle distribution. differences in dispersal can be explained qualitatively by differences in net distance travelled. These differences are also consistent with the measured velocities and tortuosities, with faster and less tortuous movement paths leading to more displacement and dispersal. We always report median and quartiles over 8 replicates. The letters indicate significant differences between species (*p <* 0.05; GLMMs and Tukey post-hoc contrasts; see S4 for details).

### Informed dispersal and movement in metacommunities

Dispersal and the underlying movement strategies were strongly impacted by the presence of competing species (Fig. 3). For all statistical analyses please refer to Tab. S5. The strength of inter-specific competition (Tab. 1) was estimated in additional experiments by fitting Lotka-Volterra models of inter-specific competition to the observed community dynamics (see Supporting Information and Fig. S2). In two-species metacommunities *Tetrahymena pyriformis* always exhibited higher dispersal rates compared to the single-species reference (Fig. 3 A and E). The presence of *Colpidium* sp., a relatively strong competitor (*α*_*Tet,Col*_ = 2.85; Tab. 1), led to a 34 % increase in the dispersal rate (Fig. 3 A). The even stronger competitor *Paramecium aurelia* (*α*_*Tet,Pau*_ = 18; Tab. 1) increased *Tetrahymena*’s dispersal rate by 780 % (Fig. 3 E). In both cases, increased dispersal rates were reflected in altered movement patterns: the presence of competitors led to higher net distances travelled (20 %, respectively 30 %; Fig. 3 B and F) due to higher movement velocities (Fig. 3 C and G). We therefore found a generally positive relationship between the strength of competitive interactions and dispersal in *Tetrahymena*. The same positive relationship held for net distance travelled as well as movement velocity and the strength of competitive interactions (Fig. S7).

**Figure 3:**
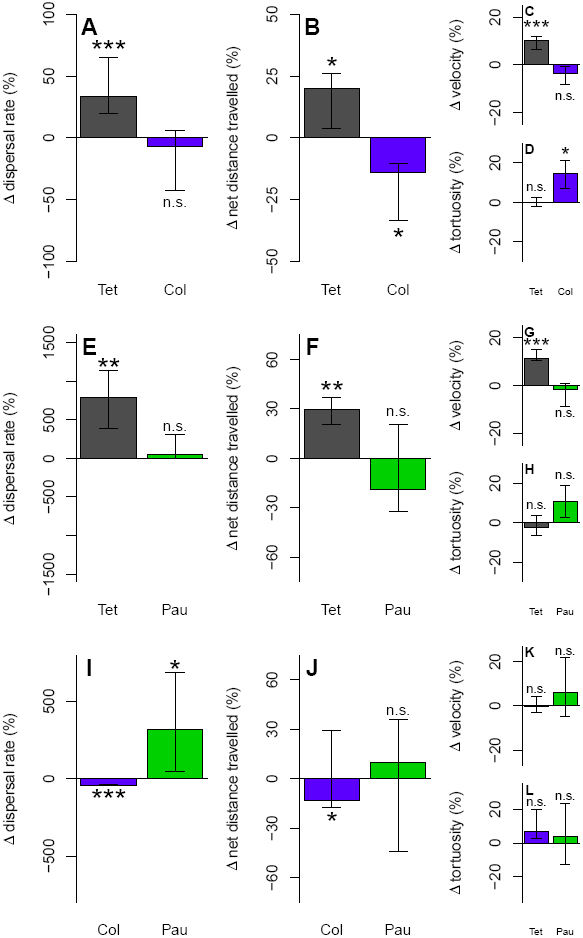
Difference in dispersal and movement behaviour between single species metapopulations and two species metacommunities. Positive (negative) values indicate more (less) dispersal/ movement in the metacommunity relative to the single species context. (A) – (D) *Tetrahymena pyriformis* (Tet) and *Colpidium* sp. (Col); (E) – (H) *Tetrahymena pyriformis* (Tet) and *Paramecium aurelia* (Pau); (I) –*Colpidium* sp. (Col) and *Paramecium aurelia* (Pau). We always report median and quartiles of the differences of the 8 replicates. The statistical analysis was performed using one-sample t-tests (*μ* = 0) on the differences (see Tab. S5). *: *p <* 0.05, **: *p <* 0.01, ***: *p <* 0.001. Note that the y-axes extensions differ among panels.

*Colpidium* sp. did not alter its dispersal rate in the presence of the weak competitor *Tetrahymena pyriformis* (*α*_*Col,Tet*_ = 0.32; Tab. 1; Fig. 3 A). On the contrary, it reduced its net distance travelled (14 %; Fig. 3 B) by increasing movement tortuosity (Fig. 3 D). The stronger competitor, *Paramecium aurelia* (*α*_*Col,Pau*_ = 2.53; Tab. 1), even decreased the dispersal rate of *Colpidium* sp. (41 %; Fig. 3 I) which was associated with a decrease in net distance travelled (13 %; Fig. 3 J) and a non-significant increase in tortuosity (Fig. 3 L).

The strongest competitor in all pairwise comparisons, *Paramecium aurelia* (*α*_*Pau,Tet*_ = 0.06, *α*_*Pau,Col*_ = 0.01; Tab. 1), did not react to the presence of *Tetrahymena pyriformis*, although a non-significant trend to smaller net distances travelled as a consequence of more tortuous paths could be observed (Fig. 3 F and H). By contrast, *Colpidium* sp., which is larger than *Tetrahymena pyriformis*, induced an important increase in dispersal rate (320 %; Fig. 3 I) which was accompanied by a non-significant increase in net distance travelled (Fig. 3 J).

### Scaling up from two to three-species metacommunities

The patterns of condition-dependent dispersal and movement observed in the two-species metacommunities (Fig. 3) were also found in the three-species metacommunity (Fig. 4). The presence of competing species induced increased dispersal in *Tetrahymena pyriformis* in comparison to single-species metapopulations (870 %; Fig. 4 A). This was associated with corresponding changes in net distance travelled and movement velocity (Fig. 4 B and C). *Colpidium* sp. was found to have a reduced dispersal rate in the three-species metacommunity (50 %; Fig. 4 A) as in the presence of *Paramecium aurelia* only (Fig. 3 I). This pattern was accompanied by a reduction in net distance travelled (Fig. 4 B) and velocity (Fig. 4 C) as well as an increase in movement tortuosity (Fig. 4 D). Similarly, *Paramecium aurelia* had an increased dispersal rate in the three-species metacommunity (327 %; Fig. 4 A), as in the two-species metacommunity with *Colpidium* sp. (Fig. 3 I). As in the two-species metacommunity this change was not strongly reflected in changes in movement due to the large amount of variation.

**Figure 4:**
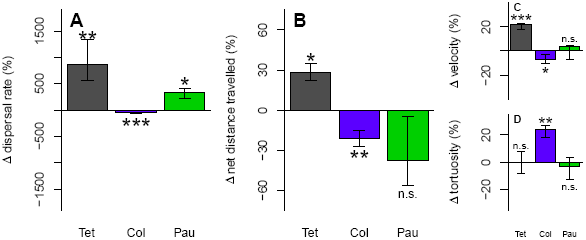
Difference in dispersal and movement behaviour between single species metapopulations and three species metacommunities (*Tetrahymena pyriformis* (Tet), *Colpidium* sp. (Col) and *Paramecium aurelia* (Pau)). Positive (negative) values indicate more (less) dispersal/ movement in the metacommunity relative to the single species context. We always report median and quartiles of the differences of the 8 replicates. The statistical analysis was performed using one-sample t-tests (*μ* = 0) on the differences (see Tab. S5). *: *p <* 0.05, **: *p <* 0.01, ***: *p <* 0.001.

## Discussion

We combined experiments in single-species metapopulations, two- and three-species metacommunities to understand the effect of interspecific competition on informed movement and dispersal. Our experiments show that the presence of competing species leads to consistent and predictable changes in dispersal and movement dynamics in experimental metacommunities (Fig. 5). This strongly indicates that dispersal and movement are informed with respect to inter-specific competition, and that inter-specific interactions have the potential to impact metacommunity dynamics and to change the number of coexisting species at local and regional scales. Furthermore, besides only depending on the strength of competitive interactions we could show that dispersal and movement also depend on the specific dispersal and movement behaviour of the interaction partners. This dependence on community composition can only be understood if one takes into account the long-term fitness-relevant consequences of interactions. Therefore, the specific multispecies context may introduce emergent phenomena that can only be understood in the metacommunity context and cannot be predicted from single-species behaviour only. As a consequence, our work suggests that predictions based on dispersal and movement estimates gained from species in isolation are likely to fail.

**Figure 5:**
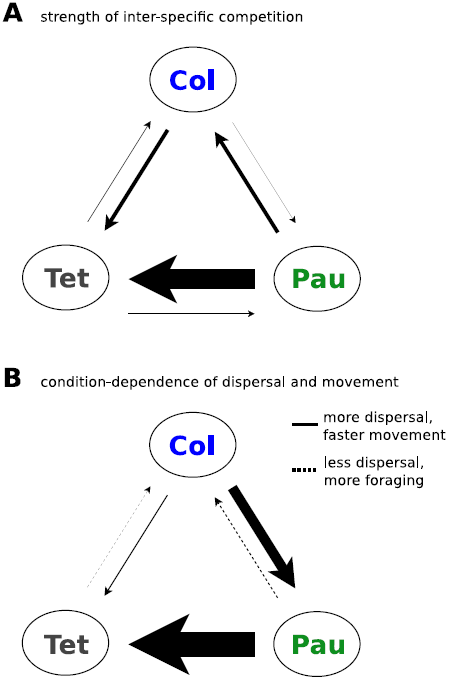
Informed movement and dispersal in metacommunities of competing species. (A) Strength of inter-specific competitive interactions in all pairwise comparisons. (B) Resulting impact on dispersal and movement behaviour. Faster and straighter movement translates (non-linearly) into more dispersal. We interpret slower and/or more tortuous movement as foraging behaviour. The width of all arrows scales with effect strength. See also Tab. 1 and Tab. S3.

### Dispersal and movement are informed with respect to inter-specific competition

Consistent with our theoretical predictions (Fig. 1), a generally positive condition-dependence of dispersal and movement was found for the least competitive species, *Tetrahymena pyriformis* (Fig. 3, 4 and Fig. S7). These results imply that models of informed dispersal, more exactly density-dependent dispersal (Metz & Gyllenberg, 2001; Poethke & Hovestadt, 2002) can be transferred from a single-species metapopulation context to a multi-species metacommunity context as indicated in Fig. 1 if one takes into account inter-specific interactions.

Interestingly, for one species, *Colpidium* sp., the effect of inter-specific competition was opposite to the results for the other species: dispersal decreased in the presence of all competitors (Fig. 3 I, 4 and Fig. S7). We suggest that this is due to a u-shaped competition-dependent dispersal/movement reaction norm depicted in Fig. 1 (Kim *et al.*, 2009; Fronhofer *et al.*, 2015). Likely, *Colpidium* sp. Shows the highest dispersal rate of all species in a single species context (Fig. 2 A) as a consequence of a relatively high species-specific Allee threshold (which we found to be true; see Fig. S5 E). In that case, an increase in competition may generally lead to a decrease in dispersal as evident from Fig. 1 given that the competition-dependent dispersal reaction norm is shifted enough to the right. Clearly, increasing competition even further should eventually lead to positive condition-dependence.

The most competitive species, *Paramecium aurelia*, which is also least affected by the presence of both other species (*α*_*Pau,j*_ is always very small; Tab. 1) did not react to increased competition in a systematic manner (Fig. S7). Such a pattern can be understood and possibly generalized if one takes the impact of the focal species (here: *P. aurelia*) on the other species into account. 1) For highly asymmetric interactions (in our case this would be between *Paramecium aurelia* and *Tetrahymena pyriformis* ; *α*_*Pau,Tet*_ = 0.06 and *α*_*Tet,Pau*_ = 18), the resulting impacts on dispersal and movement behaviour will also be highly asymmetric as the weaker competitor has nearly no fitness-relevant impact on the focal species, and we indeed found an increase in the dispersal rate of *Tetrahymena pyriformis* by 780 %. Subsequently, the weaker competitor will eventually leave the local community and the stronger competitor does not experience any fitness advantages by altering its dispersal or movement behaviour. 2) If however, the weaker competitor happens to suffer from a strong Allee effect (here: *Colpidium* sp.) its dispersal rate may be decreased by the focal species, as described above. This has important consequences as competition will increase locally and the weaker competitor, if strong enough, will start to have fitness-relevant negative effects on the focal species. While being a better competitor, the focal species could then experience fitness benefits by increasing its dispersal rate. This scenario potentially explains why *P. aurelia* consistently showed an increase in dispersal and movement in the presence of *Colpidium* sp., a weaker competitor, in both two- and three-species metacommunities (Fig. 3 I and 4). Consequently, dispersal and movement are not only informed with respect to inter-specific competition but can also depend on the specific dynamic behaviour of the interaction partner and the long-term consequences of this behaviour.

### Scaling from two- to three-species metacommunities

Our results for two- and three-species metacommunities show that informed dispersal and movement is consistent across metacommunities of increasing diversity (Fig. 3 and 4). The major patterns observed in the two-species systems, namely a general increase in dispersal and movement for *Tetrahymena pyriformis*, a decrease in dispersal and movement for *Colpidium* sp. in the presence of *Paramecium aurelia* and a simultaneous increase in dispersal for *Paramecium aurelia* (Fig. 3), can all be recaptured in the three-species metacommunity (Fig. 4). The relative changes in effect size suggest a somewhat monotonic effect. This is encouraging, as it suggests that predictions which scale up from simple to more complex communities are, in principle, possible and not elusive due to strong non-linearities.

Finally, all our experiments show that local movement behaviour can be used to explain differences in dispersal (Fig. 2 and Fig. S6). Straighter and faster movement leads to more dispersal while slower and more tortuous movement leads to less dispersal. Specifically, increased dispersal is mostly linked to higher velocities, while decreased dispersal and foraging-like behaviour is primarily linked to an increase in tortuosity (Fig. 3 and 4). This holds across all species investigated in this study, although the quantitative relationship are species-specific (Fig. S6). The relationship between movement and dispersal is non-linear, such that, at specific starting conditions, already small differences in movement can lead to large differences in dispersal (Fig. S6).

As a possibly interesting extension, our work also provides a mean of inferring dispersal rates from an interaction matrix (e.g. Tab. 1) and a reaction norm (e.g. Fig. 1). As dispersal, and the incentive to disperse, are notoriously difficult to assess in natural communities, the often much better known interaction terms (e.g. Carrara *et al.*, 2015) may be a way to infer relative changes in dispersal rates across different communities.

### Metacommunity consequences of informed dispersal and movement

While information use is increasingly taken into account in single-species metapopulations it has largely remained unexplored in a multi-species metacommunity context (but see Amarasekare 2010 for a theoretical model and De Meester *et al.* 2015 for empirical evidence). Our work demonstrates the importance of considering informed, non-random processes in studies of metacommunities, especially given the large effect sizes we report here (e.g. increases in dispersal rates up to 800 % for some species; Fig. 3 and 4). The metacommunity-level consequences of such drastic changes in dispersal remain to be explored in detail both theoretically and empirically. Given what we know about the consequences of density dependent dispersal in metapopulations (see e.g. Amarasekare, 2004; Enfjäll & Leimar, 2009; Kubisch *et al.*, 2011; Altwegg *et al.*, 2013) the consequences are likely of large importance for population synchrony and stability in natural metacommunities (Koelle & Vandermeer, 2004; Gouhier *et al.*, 2010), coexistence (Amarasekare, 2010), composition as well as the spatial distribution and temporal dynamics of food webs (Meliáan *et al.*, 2015).

It is interesting to note that we did not find the classically reported competition-colonization trade-off (e.g. Cadotte, 2007), i.e. weaker competitors showing higher dispersal abilities and vice versa, in the single species metapopulations. In Fig. 2 the species are ordered by competitive rank and the pattern is clearly unimodal. By contrast, in the multi-species metacommunities informed dispersal led to the emergence of what is usually termed “competition-colonization trade-off” as one can see in Fig. S8. Importantly, this negative correlation between dispersal and competitive ability is not a genuine genetic or cost related trade-off between dispersiveness and competitive ability. This result has potentially far-reaching implications as competition-colonization trade-offs have been linked to species coexistence and stability of metacommunites (Calcagno *et al.*, 2006; Livingston *et al.*, 2012). Such behavioural plasticity will have important consequences for metacommunity dynamics and geographic distributions which underlines the need to take informed dispersal into account.

## Conclusions

Our experimental work shows that both dispersal and movement are informed and adjusted with respect to inter-specific competition. To a large extent, models of density-dependent dispersal and movement developed for single-species metapopulations can be amended to be applicable to metacommunities, highlighting that both intra- and inter-specific interactions must be considered when understanding dispersal of species in a multi-species context. As we show that dispersal and movement strongly depend on the community context and the behavioural response of interacting species our work also implies that predictions based on dispersal and movement rates estimated in a single-species context only are likely erroneous and miss emergent, community-level phenomena.

## Acknowledgements

We would like to thank Pravin Ganesanandamoorthy for help with data collection and Regula Illi for help during laboratory work. Funding is from Eawag (EAF), Sciex Fellowship 12.327 (JK) and the Swiss National Science Foundation Grants PP00P3 150698 (FA) and 31003A 144162 (CM).

## Supporting Information

### Estimating growth rates and carrying capacities

In order to quantify growth rates (*r*_0_) and carrying capacities (*K*) of the three protist species (*Tetrahymena pyriformis* (Tet), *Colpidium* sp. (Col) and *Paramecium aurelia* (Pau)) we performed additional experiments using the same microcosms as in the main experiment (20 mL Starstedt vials, with 15 mL bacterized medium). As described in the main text, batch cultures of all species were grown to carrying capacity before the experiments. Growth curves were started by dilution at 0.1*K* and followed over 10 days (Fig. S1). Data on population densities were collected as described in the main text using video analysis (due to technical reasons videos were recorded with a Canon 5D Mark III camera; all analyses were adjusted appropriately). These experiments were replicated 6 times.

To estimate growth parameters we fit logistic growth curves to the individual replicates:

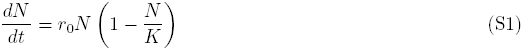

where *N* is the population size at time *t*, *r*_0_ is the growth rate and *K* the carrying capacity.

To fit the model and estimate *r*_0_ and *K* we used a least-squares approach. The differential equation was solved using the software package ****R version 3.1.2 (R Core Team, 2014) (function “ode” of the “deSolve” package, version 1.10-9). The model was fit to the measured population density data using the Levenberg-Marquardt algorithm (function “nls.lm” of the “minpack.lm” package, version 1.1-8). For results see Fig. S1 and Tab. S1.

### Estimating competition coefficients

Competition coefficients (*α*_*i,j*_) which capture the relative competitive effect of one species (*j*) on another (*i*) were estimated analogously to growth rates and carrying capacities. In additional experiments using the same microcosm set-up we recorded time-series of population densities in all two-species communities (Tet–Col, Tet–Pau and Col–Pau) using video analysis as described above and in the main text (Fig. S2). Communities were started from batch cultures that had been grown to carrying capacity. Species were added at 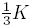 and experiments were replicated 6 times.

To estimate the pairwise competition coefficients we fit Lotka-Volterra models of inter-specific competition to the individual replicates:

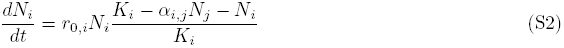

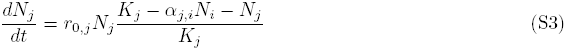

the values for growth rates (*r*_0_) and carrying capacities (*K*) were collected independently (see above). *α*_*i,j*_ represents the relative effect of species *j* on *i*. For example, if *α*_*i,j*_ = 2 species *j* has twice the competitive effect on species *i* as species *i* has on itself, *α*_*i,j*_ = 1 implies that inter- and intra-specific competition are equivalently strong and *α*_*i,j*_ = 0 suggests that species *j* does not compete with species *i*.

The fitting procedure followed a least-squares approach as described above. Since population densities varied widely (see Fig. S1 and Tab. S1) the fitting procedure used data normalized to carrying capacity. As the recorded community dynamics did not indicate facilitation (the measured densities did not consistently increase to values over carrying capacity) we set the fitting algorithm’s lower boundary for possible *α*_*i,j*_ values to zero, i.e. no interaction. After preliminary analyses and optical inspection of the fits the upper boundary was set to 18 in order to allow for timely convergence. This boundary only impacts the values estimated for *α*_*Tet,Pau*_ which otherwise increase to extremely higher values. See Fig. S2 for data and fits. See Tab. S3 for fitted values.

### Correction of dispersal rates

Due to large differences in dispersal ability (see Fig. 2) of our three study species the dispersal time window we used for the experiment was relatively long (14 h). Importantly, dispersal and growth could be confounded, as the generation time of the fastest reproducing species (Tet, see Fig. S1 and Tab. S1) is well below the length of the experiment. In order to avoid this bias we used two-patch metapopulation models to estimate the growth-corrected dispersal rates (*d*_*corr*_) based on the observed population densities at the end of the dispersal phase in start and target patches. In a first step we assume that populations in both, start and target patch, grow logistically and that dispersal is symmetric and free of costs:

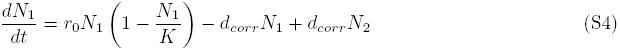

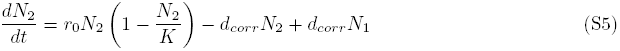

where *N*_1_ is the population size in the start patch, *N*_2_ the population size in the target patch, *r*_0_ is the growth rate and *K* the carrying capacity. Both *N*_1_ and *N*_2_ were measured at the beginning (*t* = 0*h*: *N*_1_ *≈* 0.25*K*, *N*_2_ = 0) and at the end of the experiment (*t* = 14*h*). *r*_0_ and *K* were estimated independently (see Fig. S1 and Tab. S1).

As we describe in the main text, there is ample evidence that suggests that our study species suffer from Allee effects (Christensen *et al.*, 2001; Chaine *et al.*, 2010; Fronhofer *et al.*, 2015; Duncan *et al.*, 2011; Odum & Allee, 1954). Allee effects, reduced growth at low densities (see Fig. S4), are highly relevant for metapopulation and -community dynamics, especially if dispersal occurs into empty patches, as in our experiments. Therefore, the presence of an Allee effect may additionally bias the directly measured dispersal rates. In order to explore whether Allee effects are relevant in our system and to avoid additional bias in the growth-corrected dispersal rates (*d*_*corr,Allee*_) we included a two-patch metapopulation model with an Allee effect into our analysis:

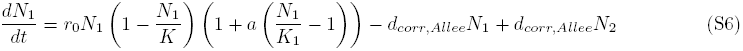

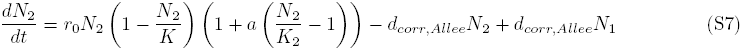

The Allee effect strength *a* increases from 0, i.e. no Allee effect, and is positive (see Fig. S4). Note that *a* = 1 is equivalent to setting the Allee threshold to *A* = 0 in the model provided by Amarasekare (1998). To fit the models and estimate *d*_*corr*_, respectively *d*_*corr,Allee*_ and *a*, we used a least-squares approach as described above.

A formal comparison of the models using the AIC criterion (Tab. S2) revealed that the model including the Allee effect (Eqns. S6 and S7) explained the data better than the model without (Eqns. S4 and S5). This implies that all three species suffer from relatively strong Allee effects, as one can see in Fig. S5. In addition, we show that, the measured dispersal rates correlate highly with the growth-corrected estimates (Tab. S2; LM: *N* = 23, *t* = 40.58, *p <* 0.001, *R*^2^ = 0.987; Fig. S5). Therefore, we performed all analyses using the originally collected and uncorrected dispersal data. See Tab. S2 for estimates of *d*_*corr*_, *d*_*corr,Allee*_ and *a* and a formal comparison with a two-patch model ignoring the Allee effect.

## Supplementary figures

**Supplementary Figure S1:**
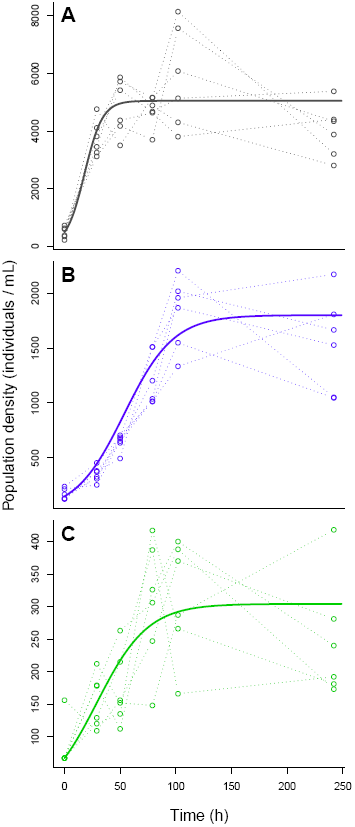
Growth curves of (A) *Tetrahymena pyriformis*, (B) *Colpidium* sp. and (C) *Paramecium aurelia* used to estimate growth rates (*r*_0_) and carrying capacities (*K*). Open circles and dotted lines represent data of the 6 independent replicates and continuous lines model predictions (logistic growth model; Eq. S1) using the median of the per replicate estimated growth parameters. See also Tab. S1.

**Supplementary Figure S2:**
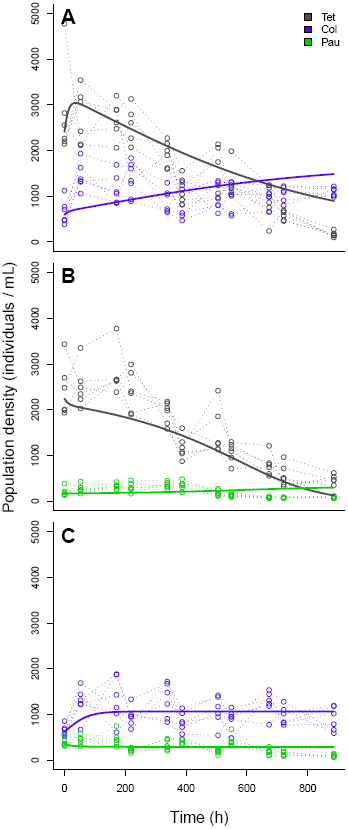
Community dynamics in all two-species communities used for the estimation of pairwise competition coefficients (*α*_*i,j*_). (A) *Tetrahymena pyriformis* (Tet) and *Colpidium* sp. (Col), (B) *Tetrahymena pyriformis* (Tet) and *Paramecium aurelia* (Pau) and (C) *Colpidium* sp. (Col) and *Paramecium aurelia* (Pau). Open circles and dotted lines represent data of the 6 independently replicated communities and continuous lines model predictions (Lotka-Volterra model of inter-specific competition; Eqns. S2 and S3) using the median of the per replicate estimated pairwise interaction coefficients and the growth rates estimated in Fig. S1. See also Tab. S3.

**Supplementary Figure S3:**
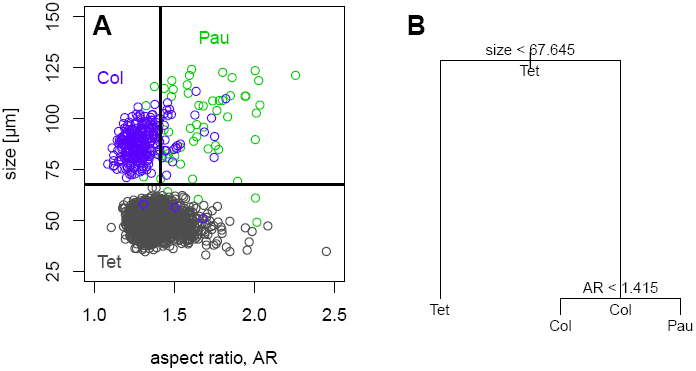
Classification of species in communities using a classification tree (Statistical Software Package R version 3.1.2; package “tree” version 1.0-35, function “tree”) with aspect ratio (AR) and length along the major body axis (size) as explanatory variables. Misclassification error rate: 0.0197.

**Supplementary Figure S4:**
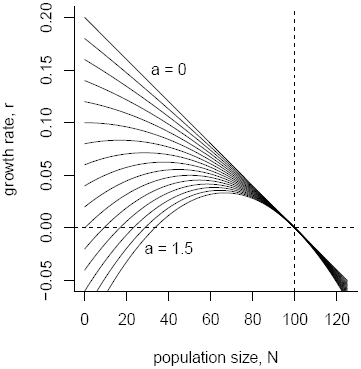
Logistic growth including an Allee effect (see Eqns. S6 and S7). Increasing *a* increases the Allee effect strength. Here depicted from *a* = 0 to *a* = 1.5 in steps of 0.1.

**Supplementary Figure S5:**
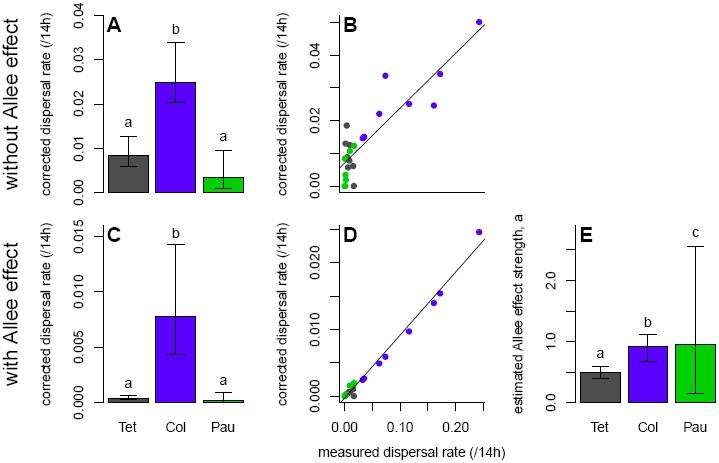
S(A) Corrected dispersal rates using a logistic growth model without (A–B) respectively with an Allee effect (C–E). See Tab. S2 for more details. See Fig. 2 for the analogous plot of the measured data. (A) For corrected data estimated from the model without an Allee effect (see Eqns. S4 and S5) the dispersal rate of *Colpidium* sp. is significantly higher than the rates of the other two species. GLMM: *N* = 23, *t*(*Pau*) = 3.47, *p*(*Pau*) = 0.0005, *t*(*Tet*) = 3.43, *p*(*Tet*) = 0.0005 (random *-*effect: replicate; error family: gamma). Post-hoc Tukey contrasts: Tet–Col: *z* = 3.43, *p* = 0.0014, Col–Pau: *z* = 3.47, *p* = 0.0015, Tet–Pau: *z* = −1.2, *p* = 0.44. (B) Correlation of corrected and measured dispersal rates (LM: *N* = 23, *t* = 8.62, *p <* 0.001, *R*^2^ = 0.769). The solid line is the model fit. (C) For corrected data estimated from the model with an Allee effect (see Eqns. S6 and S7) the dispersal rate of *Colpidium* sp. is significantly higher than the rates of the other two species. GLMM: *N* = 23, *t*(*Pau*) = 2.73, *p*(*Pau*) = 0.006, *t*(*Tet*) = 4.46, *p*(*Tet*) *<* 0.001 (random effect: replicate; error family: gamma). Post-hoc Tukey contrasts: Tet–Col: *z* = 4.46, *p <* 0.001, Col–Pau: *z* = 2.73, *p* = 0.016, Tet–Pau: *z* = 1.14, *p* = 0.46. (D) Correlation of corrected and measured dispersal rates (LM: *N* = 23, *t* = 40.58, *p <* 0.001, *R*^2^ = 0.987). The solid line is the model fit. (E) Estimated Allee effect strengths (*a*). GLMM: *N* = 23, *t*(*Pau*) = − 3.19, *p*(*Pau*) = 0.0014, *t*(*Tet*) = 2.82, *p*(*Tet*) = 0.0049 (random effect: replicate; error family: gamma). Post-hoc Tukey contrasts: Tet–Col: *z* = 2.82, *p* = 0.013, Col–Pau: *z* = −3.19, *p* = 0.0036, Tet–Pau: *z* = 4.89, *p <* 0.001. We always report median and quartiles over the 8 replicates.

**Supplementary Figure S6:**
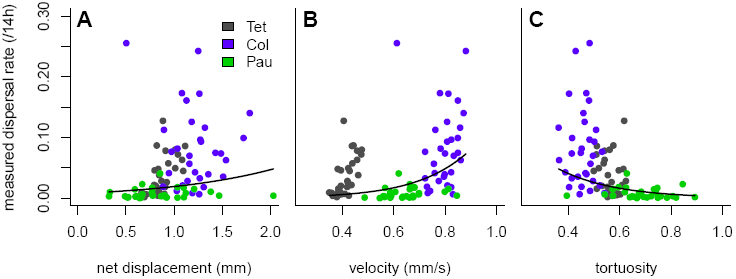
Correlation between dispersal and movement across all species in all metapopulations and -communities. The strong non-linearity explains why small differences in displacement led to large differences in dispersal in the data. (A) Dispersal as a function of the net distance travelled (GLMM: *N* = 90, *z* = 1.99, *p* = 0.046; random effects: replicate and species; observation-level random effect due to overdispersion; error family: binomial). (B) Dispersal as a function of velocity (GLMM: *N* = 90, *z* = 2.83, *p* = 0.005; random effects: replicate and species; observation-level random effect due to overdispersion; error family: binomial). (C) Dispersal as a function of tortuosity (GLMM: *N* = 90, *z* = −2.09, *p* = 0.037; random effects: replicate and species; observation-level random effect due to overdispersion; error family: binomial).

**Supplementary Figure S7:**
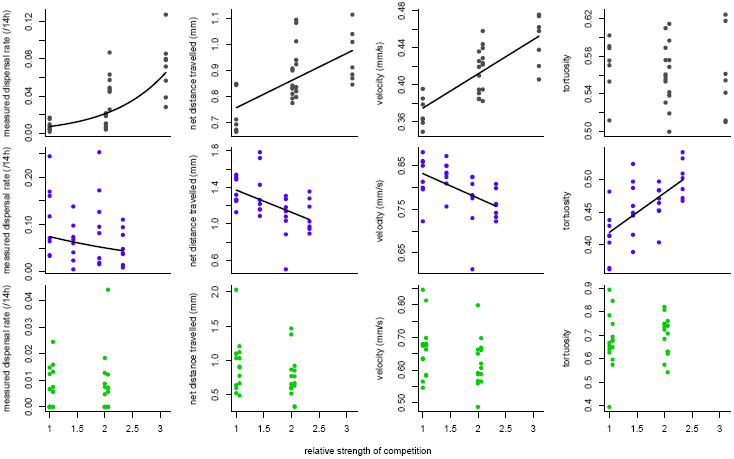
Dispersal and movement parameters as a function of the relative strength of competition in all metacommunties. The relative strength of competition follows the logic of Fig. 1 and is calculated as the sum of all individuals in the starting patch across species at the beginning of the experiment weighted by the specific competition coefficients relative to the single species density (see Tab. S3). Therefore the relative strength of competition in every single species metapopulation is by definition 1. In a Tet–Col metacommunity the relative strength of competition experienced by Col is calculated as: (0.25*K*_*Col*_ + 0.25*K*_*Tet*_*α_Col,Tet_*)/(0.25*K*_*Col*_), since all experiments were started with densities at 0.25*K*. Regression lines are significant fixed effects of GLMM respectively LMM fits (see Tab. S6 for the statistical analysis). Values from all replicates are shown.

**Supplementary Figure S8:**
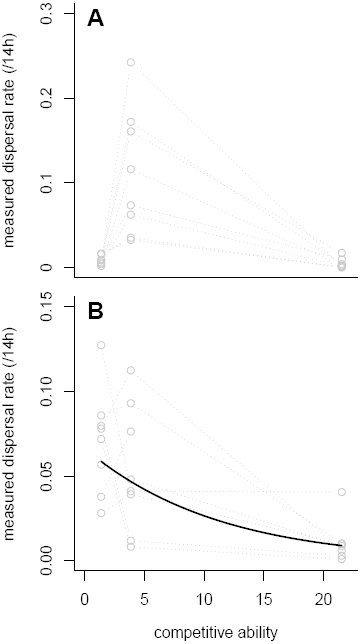
Dispersal as a function of competitive ability across all three study species. Competitive ability is calculated as the row sums of Tab. S3. (A) Dispersal rates in the single species metapopulations. For a detailed analysis see Fig. 2. (B) Dispersal rates in the three-species metacommunity (GLMM: *N* = 23, *z* = −4.67, *p <* 0.001; random effects: replicate; observation-level random effect due to overdispersion; error family: binomial). Clearly, a “competition-colonization trade-off” (e.g. Calcagno *et al.*, 2006; Cadotte *et al.*, 2006; Cadotte, 2007; Livingston *et al.*, 2012) can only be observed here and is the result of condition-dependent dispersal/ movement behaviour. Consequently, our results suggest that a genuine, intra-specific trade-off due to costly dispersal and competitive behaviour does not exists, rather a negative correlation emerges only in the multi-species system due to behavioural plasticity.

## Supplementary tables

**Supplementary Table S1:** Density-dependent growth in single species populations. Parameters were estimated by fitting a logistic growth model (Eqn. S1) to each of 8 replicates individually using a least squares approach (Levenberg-Marquardt algorithm). The reported values are medians and the first and third quartile in brackets. *r*_0_ is given in *h*^−1^ and *K* in individuals per mL. See also Fig. S1.

|  | *T. pyriformis* (Tet) | *Colpidium sp.* (Col) | *P. aurelia* (Pau) |
| --- | --- | --- | --- |
| $r_0$ | 0.12 (0.1, 0.15) | 0.05 (0.04, 0.05) | 0.04 (0.03, 0.06) |
| $K$ | 5047 (4570, 5144) | 1800 (1653, 1858) | 304 (277, 321) |

**Supplementary Table S2:** Corrected dispersal rates estimated from two-patch metapopulation models with and without an Allee effect (see Eqns. S4 – S7). A formal comparison of the two models using AIC indicates that the model including the Allee effect is more appropriate for our study species. All dispersal rates are in %. The missing values in the third replicate of Pau are due to missing counts of starting conditions. See also Fig. S5.

| species | replicate | $d_{obs}$ | $d_{corr}$ | $d_{corr, Allee}$ | $a$ | $AIC(d_{corr})$ | $AIC(d_{corr, Allee})$ |
| --- | --- | --- | --- | --- | --- | --- | --- |
| Tet | 1 | 1.6674 | 0 | 0 | 0 | 29.10 | 31.10 |
| Tet | 2 | 1.5502 | 0.6097 | 0.1031 | 0.4018 | 35.50 | -91.98 |
| Tet | 3 | 0.8447 | 0.7778 | 0.0584 | 0.4823 | 36.26 | -88.76 |
| Tet | 4 | 0.4841 | 0.8837 | 0.0329 | 0.4023 | 35.40 | -99.76 |
| Tet | 5 | 0.6140 | 0.5669 | 0.0458 | 0.6936 | 37.33 | -98.61 |
| Tet | 6 | 0.1787 | 1.2970 | 0.0128 | 0.5470 | 36.48 | -129.11 |
| Tet | 7 | 0.9247 | 1.2546 | 0.0659 | 0.5181 | 36.31 | -101.39 |
| Tet | 8 | 0.3868 | 1.8504 | 0.0307 | 0.9194 | 38.65 | -94.57 |
| Col | 1 | 3.2634 | 1.4597 | 0.2425 | 0.6292 | 26.27 | -105.84 |
| Col | 2 | 3.5309 | 1.5049 | 0.2644 | 0.6868 | 27.31 | -104.46 |
| Col | 3 | 17.2088 | 3.4238 | 1.5402 | 1.0162 | 28.43 | -100.92 |
| Col | 4 | 24.2321 | 5.0168 | 2.4603 | 1.4870 | 30.24 | -98.14 |
| Col | 5 | 16.0784 | 2.4630 | 1.3959 | 0.6458 | 27.69 | -94.36 |
| Col | 6 | 11.6110 | 2.5117 | 0.9691 | 0.9229 | 29.00 | -109.89 |
| Col | 7 | 7.3529 | 3.3736 | 0.5894 | 1.4152 | 29.05 | -112.89 |
| Col | 8 | 6.2500 | 2.2082 | 0.4875 | 0.9187 | 28.07 | -102.59 |
| Pau | 1 | 0.0764 | 0.8359 | 0.0057 | 0.9498 | 22.26 | -99.76 |
| Pau | 2 | 0 | 0 | 0 | 0 | 14.06 | 16.06 |
| Pau | 3 | 0.4024 | – | – | – | – | – |
| Pau | 4 | 0.1387 | 0 | 0 | 0 | 24.8259 | 26.8259 |
| Pau | 5 | 1.7241 | 1.2296 | 0.1950 | 3.0876 | 26.31 | -106.03 |
| Pau | 6 | 0.2445 | 0.1877 | 0.0165 | 0.3051 | 17.57 | -123.32 |
| Pau | 7 | 0.9492 | 1.0698 | 0.1591 | 5.1494 | 26.94 | -109.70 |
| Pau | 8 | 0.1664 | 0.3400 | 0.0161 | 2.0416 | 22.59 | -108.08 |

**Supplementary Table S3:** Interaction coefficients (*α*_*i,j*_) of a Lotka-Volterra model of inter-specific competition (see Eqns. S2 and S3). Species *i* can be found in the columns and species *j* in the rows, so that *α*_*Tet,Col*_ = 2.85, for example. Reported values are median values and in brackets quartiles of 8 independently fitted replicates. For the metacommunity dynamics and model fits see Fig. S2. *: Variation is zero here as the values represent maximal parameter values imposed on the fitting algorithm. See text for further explanations.

|  | Tet | Col | Pau |
| --- | --- | --- | --- |
| Tet | 1 | 0.32 (0.26, 0.67) | 0.06 (0.053, 0.21) |
| Col | 2.85 (2.68, 14.2) | 1 | 0.01 (0.001, 0.02) |
| Pau | 18 (18, 18)* | 2.53 (2.20, 2.74) | 1 |

**Supplementary Table S4:** Statistical analysis of dispersal and movement behaviour in single-species metapopulations for all three study species (*Tetrahymena pyriformis* (Tet), *Colpidium* sp. (Col) and *Paramecium aurelia* (Pau)). See Fig. 2 for a graphical representation.

|  | N | generalized linear mixed model |  |  |  | random effects | error family | post-hoc Tukey contrasts |  |  |  |
| --- | --- | --- | --- | --- | --- | --- | --- | --- | --- | --- | --- |
|  |  | Col (intercept) | Pau | Tet |  |  |  | Tet-Col | Col-Pau | Tet-Pau |  |
| dispersal rate | 24 | $z = -10.05, p < 0.001$ | $z = -5.85, p < 0.001$ | $z = -7.61, p < 0.001$ | | rep., obs. | binomial | $z = -7.61, p < 0.001$ | $z = -5.85, p < 0.001$ | $z = 1.27, p = 0.41$ | |
| net dist. travelled | 2189 | $t = 17.80, p < 0.001$ | $t = 3.09, p = 0.002$ | $t = 13.86, p < 0.001$ | | rep. | gamma | $z = 18.86, p < 0.001$ | $z = 3.09, p = 0.004$ | $z = 1.63, p = 0.23$ | |
| velocity | 2189 | $t = 30.99, p < 0.001$ | $t = 8.52, p < 0.001$ | $t = 107.16, p < 0.001$ | | rep. | gamma | $z = 107.16, p < 0.001$ | $z = 8.52, p < 0.001$ | $z = 37.8, p < 0.001$ | |
| tortuosity | 2189 | $t = 42.54, p < 0.001$ | $t = -12.31, p < 0.001$ | $t = 13.61, p < 0.001$ | | rep. | gamma | $z = -13.61, p < 0.001$ | $z = -12.31, p < 0.001$ | $z = 4.84, p < 0.001$ | |

**Supplementary Table S5:**
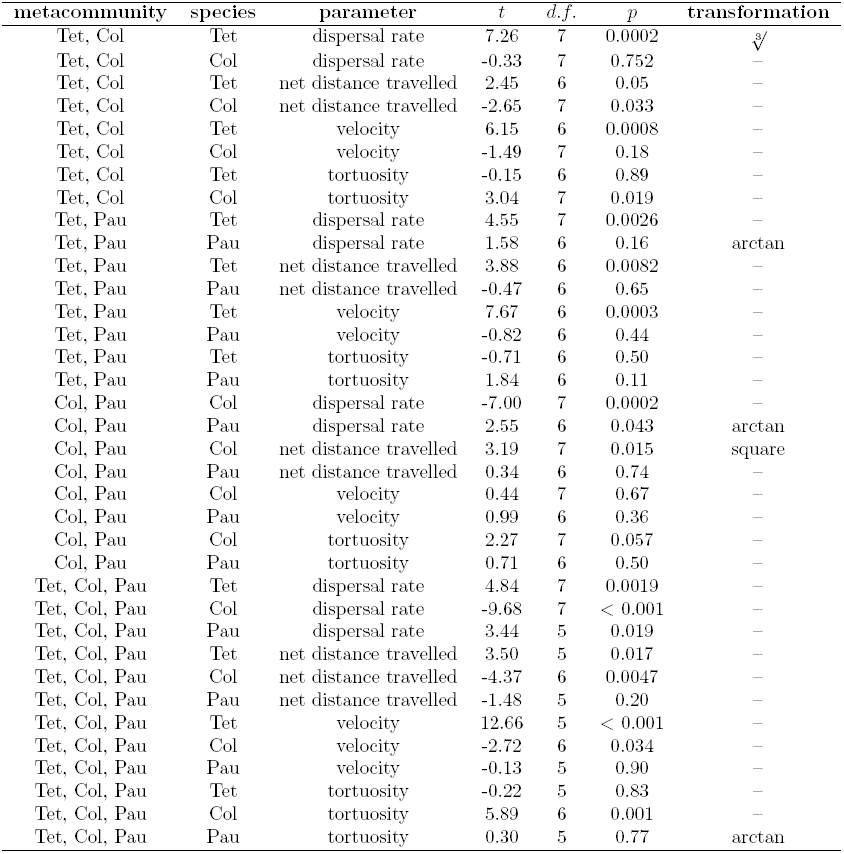
Statistical analysis of the difference in dispersal and movement between single species metapopulations and multi-species metacommunities (see Figs. 3 and 4). The analysis was done using one-sample t-tests (*μ* = 0) on the differences. When appropriate, data were transformed as indicated.

**Supplementary Table S6:** Statistical analysis of the correlation between dispersal, respectively movement, and the relative strength of competition in the two-species metacommunities (see Fig. S7). The analysis was done using LMMs and GLMMs where appropriate. Note that the test statistic reported is a *t*-statistic for LMMs and a *z*-statistic for GLMMs. Observation level random effects are used in case of overdispersion.

| **species** | **response variable** | **$N$** | **$t$ or $z$** | **$p$** | **random effect(s)** | **error family** |
| --- | --- | --- | --- | --- | --- | --- |
| Tet | dispersal rate | 32 | 6.42 | < 0.001 | replicate, observation | binomial |
| Tet | net distance travelled | 30 | 4.60 | 0.0001 | replicate | gaussian |
| Tet | velocity | 30 | 10.83 | < 0.001 | replicate | gaussian |
| Tet | tortuosity | 30 | -0.45 | 0.66 | replicate | gaussian |
| Col | dispersal rate | 32 | -3.16 | 0.0016 | replicate | binomial |
| Col | net distance travelled | 31 | -2.95 | 0.0063 | replicate | gaussian |
| Col | velocity | 31 | -3.07 | 0.0046 | replicate | gaussian |
| Col | tortuosity | 31 | 4.78 | < 0.001 | replicate | gaussian |
| Pau | dispersal rate | 31 | 0.78 | 0.44 | replicate | binomial |
| Pau | net distance travelled | 30 | -1.72 | 0.10 | replicate | gaussian |
| Pau | velocity | 30 | -1.47 | 0.16 | replicate | gaussian |
| Pau | tortuosity | 30 | 0.58 | 0.57 | replicate | gaussian |

